# Decreased metabolism and increased tolerability to extreme environment in *Staphylococcus warneri* with long-term space flight

**DOI:** 10.1101/501163

**Authors:** Po Bai, Bin Zhang, Xian Zhao, Diangeng Li, Yi Yu, Xuelin Zhang, Bing Huang, Changting Liu

## Abstract

**Background:** Many studies have shown that space environment can affect bacteria to cause a range of mutations. However, so far, there are less studies on the effects of long-term space flight (>1 month) on bacteria. In this study, a *Staphylococcus warneri* strain, which was isolated from the Shenzhou-10 spacecraft that had experienced a space flight (15-day), was carried into space again. After 64-day flight, the combined phenotypic, genomic, transcriptomic and proteomic analyses were performed to compare the influence of the two space flights on this bacteria.

**Results:** Compared with short-term space flight, long-term space flight increased the biofilm formation ability of *S. warneri* and the cell wall resistance to external environment stress, but reduced the sensitivity to chemical stimulation. Further analysis showed that these changes might be related to the significantly up-regulated gene expression of phosphotransferase system, which regulated glucose metabolism pathway, including glucose, mannose, fructose and cellobiose. The mutation of *S. warneri* caused by 15-day space flight was limited at phenotype and gene level after ground culture.

**Conclusion:** After a 79-day space flight, the changes of *S. warneri* are meaningful. Phosphotransferase system of *S. warneri* was up-regulated by long-term space stimulation, which resulted in a series of changes in cell wall, biofilm and chemical sensitivity, thus enhancing the resistance and adaptability of bacteria to the external environment.

**Importance:** Taking *S. warneri* as an example, we found that the mutation of bacteria in long-term space flight mainly focused on the adaptation and tolerance like environmental adaptation, physical resistance and cell repair to various pressures and these changes should be more obvious or even become stable mutations. But the clinical changes, for example drug resistance and virulence should not be significant because there is no threat of antibiotics.

## 1. Introduction

*Staphylococcus warneri* is a member of coagulase-negative *Staphylococci* (CNS). Generally, CNS species are considered to be harmless as skin commensal, but they are opportunistic pathogens and cause bacteremia and septicemia when people are immuno-compromised. It has been reported that the detection rate of CNS has exceeded *S. aureus* in the bloodstream infection **^[1]^**. For example, *S. warneri* can cause arterial embolism in infective endocarditis.

Space environment is associated with a variety of special stressors, such as low gravity, strong radiation such as ultraviolet radiation and heavy–ion radiation, and weak magnetic field, which can accelerate microbial mutation rate and lead to many varied phenotypes**^[2]^**. It is found that *Staphylococci* was the largest number of bacteria isolated from the Mir space station**^[3]^**. During spaceflight *Staphylococci* were carried into extraterrestrial environment by the space crew members. In the stressful airspace environment they may develop unexpected trait that could be important to spaceflight mission and have potential threat to general public health. Previous study has found that the expression of antibiotic resistance gene of *S. epidermidis* after space flight was up-regulated, and the mutation frequency increased obviously**^[4]^**. These uncertain variations increase the potential risk of pathogenic CNS to human, and have became a hotspot in the study of space microbiology. Like other CNS, *S. warneri* is easy to attach to artificial equipment or condensate water pipeline, which increases the chance to infect astronauts. The above effects together with the space environment and spacecraft condition could contribute to detrimental alteration in human immune system**^[5]^**. Thus, the risk caused by *S. warneri* may be bigger in space than that on the ground. In addition, so far there is no research to distinguish the influence of short and long space flight time on microorganisms. Compared to a short space flight, a long-period flight increases the exposure doses of the bacteria to the microgravity and space radiation environment, which may lead to unknown effects on bacteria’s phenotype and gene expression. It is unclear if the microgravity or the elevated radiation doses during a long space flight may or may not cause a higher mutation rate of *S. warneri* and whether this mutation is stable or not. In general, one would expect that these bacteria may reduce their metabolism activities in order to adapt and survive in the severe space environment for a long flight. However, all these questions are unknown so far. In this study, an *S. warneri* isolate was carried by Shenzhou-10, 11 spacecraft and Tiangong-2 space lab. This provided us a unique opportunity to explore the underlying mechanism about bacterial tolerability to extreme environment. Correspondingly, the phenotypic, genomic, transcriptomic and proteomic analyses were combined to examine the influence of the long-term space flight on this bacteria.

## 2. Methods

### 2.1 Bacterial strains and culture conditions

The ancestral *S. warneri* strain (designed as SWO) was obtained from the condensate water inside Shenzhou-10 spacecraft (launched in June 11, 2013 and had a 15-day spaceflight) and stored at -80°C with 20% glycerinum. Prior to the departure of Tiangong-2 spacelab, SWO was cultivated into two cryogenic vials filled with semi-solid Luria–Bertani (LB) medium, and placed under the following condition: subculture on ground as control group (designed as SWG), and subculture in Tiangong-2 spacelab launched on September 15, 2016 and returned to Earth by Shenzhou-11 spacecraft on November 18, 2016 (SWS). The rest of SWO was stored at -80°C. The SWS was placed inside the spacecraft and except space environment, all other culture conditions such as temperature, humidity level, oxygen level were identical between the two groups. After 64-day flight the return module of the spacecraft landed on earth, and then SWS was immediately transported back to the laboratory in dry ice containers (the same disposition to SWG). The three strains were resuscitated after SWS arriving at the laboratory and a series of assays were performed to explore the effect of a space environment on *S. warneri*, including morphology, biophysical feature, growth rate genomics, transcriptomics and proteomics. For transcriptomic and proteomic analysis, we sequenced SWG and SWS strains three times, respectively.

### 2.2 Phenotypic characteristics

#### (1) Scanning electron microscopy

The *S. warneri* strains, grown in LB medium, were washed three times with PBS (pH 7.4), and fixed with 4% glutaraldehyde overnight. After washed three times again with PBS (pH 7.4), the three samples were dehydrated in increasing grades of ethanol and critical point-dried. After coated with gold-palladium, the specimens were examined with a FEI Quanta 200 SEM scanning electron microscope (FEI, OR, USA).

#### (2) Drug susceptibility testing

Drug susceptibility testing (DST) was performed using the disk diffusion method. In brief, following the CLSI M100-S24 document**^[6]^** 12 antibiotics were selected to test the resistance of the three *S. warneri* strains, including ertapenem (ETP), aztreonam (ATM), ciprofloxacin (CIP), linezolid (LZD), tobramycin (TOB), trimethoprimsulfa--methoxazole (SXT), amikacin (AK), cefazolin (KZ), cefepime (FEP), vancomycin (VA), ceftriaxone (CRO), ampicillin/sulbactam (SAM). The entire surface of the Petri dish containing LB was covered with the required inoculum (10^7^–10^8^ CFU /ml), and the plate was dried for 15 min before the disks were placed on the surface. Following incubation for 18 h at 37°C, the zone diameters were measured according to the standard protocol.

#### (3) Carbon source utilization and chemical sensitivity assay

We performed carbon source utilization and chemical sensitivity tests using the 96-hole Biolog GENIII MicroPlate, which included 71 carbon source and 23 chemical compounds. Briefly, the bacterial culture were picked up from the surface of the BUG1B agar plate (Biolog, CA, USA) using a sterile cotton-tipped swab and inoculated into the IF-A Inoculum (Biolog, CA, USA). The target cell density of inoculum was set to 90-98% by turbidimeter (BioMe’rieux, Lyon, Fance). Then, 100 ml of the inoculums was added into each hole of the 96 GEN III MicroPlate_TM_ (Biolog, CA, USA). After incubating for 24 h at 37°C, OD_630_ of each hole was detected with a Biolog microplate reader automatically.

#### (4) Growth rate

Bacterial growth was monitored using the Bioscreen C system and Biolink software (Lab Systems, Helsinki, Finland). The *S. warneri* strains were grown overnight in LB liquid medium at 37°C. A 20 ul aliquot of overnight culture with aconcentration of 10^6^ CFU/ml was cultivated in Bioscreen C (Lab Systems, Helsinki, Finland) in 200-hole microtiter plates (honeycomb plates). The plates were continuously shaken at the maximum amplitude and incubated for 24 h at 37°C with 350 ul of fresh LB growth medium per hole. OD_630_ was measured each 2 hour. A blank hole with 370 ul of LB was also included. All experiments were repeated three times.

#### (5) Biofilm assay

After grown in LB liquid media overnight, 1 ul culture was added into 99 ul LB liquid media in a 96-well plate, and incubated at 30°C for 10 h. Then non-attached bacteria cells were removed and the plate was rinsed three times. One hundred twenty milliliter 0.1% crystal violet solution was added, and after rinsed three times 200 ul 95% ethanol solution was added to dissolve all the crystal violet on the wall of plates. Finally, after removing 125 ul solution from each well to a clean polystyrene microtiter dish, OD_560_ was determined in a microplate reader. Each sample was repeated three times.

### 2.3 Genome sequencing

#### (1) Genome sequencing and assembly

SWO genome was sequenced using a PacBio RS II platform and Illumina HiSeq 4000 platform at the Beijing Genomics Institute (BGI, Shenzhen, China). Four SMRT cells Zero-Mode Waveguide arrays of sequencing were used by the PacBio platform to generate the subreads set. PacBio subreads with length < 1 kb were removed. The program Pbdagcon was used for self correction. Draft genomic unities, which were uncontested groups of fragments, were assembled using the Celera Assembler against a highquality corrected circular consensus sequence subreads set. To improve the accuracy of the genome sequences, GATK and SOAP tool packages (SOAP2, SOAPsnp, and SOAPindel) were used to make single-base correction. To search the presence of any plasmid, the filtered Illumina reads were mapped using SOAP to the bacterial plasmid database.

#### (2) Genome Component prediction

Gene prediction was performed on SWO genome assembly by Glimmer3 with Hidden Markov models. tRNA, rRNA and sRNAs recognition made use of tRNAscan-SE, RNAmmer and the Rfam database, respectively. The tandem repeats annotation was obtained using the Tandem Repeat Finder and the minisatellite DNA and microsatellite DNA selected based on the number and length of repeat units. The Genomic Island Suite of Tools (GIST) used for genomic lands analysis with IslandPath-DIOMB, SIGI-HMM, and IslandPicker method. Prophage regions were predicted using the PHAge Search Tools (PHAST) web server and CRISPR identification using CRISPRFinder.

#### (3) Gene annotation and protein classification

The best hit was obtained using Blast alignment tool for function annotation. Seven databases, including Kyoto Encyclopedia of Genes and Genomes (KEGG), Clusters of Orthologous Groups (COG), Non-Redundant Protein Database databases (NR), Swiss-Prot**^[8]^**, Gene Ontology (GO), TrEMBL, and EggNOG, were used for general function annotation. For pathogenicity and drug resistance analysis, virulence factors and resistance gene were identified based on the core dataset in Virulence Factors of Pathogenic Bacteria (VFDB), Antibiotic Resistance Genes Database (ARDB), Pathogen Host Interactions (PHI) and Carbohydrate-Active Enzymes Database (CAED). Type III secretion system effector proteins were detected by EffectiveT3.

### 2.4 Comparative genomic analysis

#### (1) SNP detection

With alignment software MUMmer**^[9]^**, each query sequence from SWS and SWG was aligned with the reference sequence. The variation sites between the query sequence and reference sequence were found out and filtered preliminarily to detect potential SNP sites. The sequences with the length of 100 bp at both sides of SNP in the reference sequence were extracted and aligned with assembly result to verify SNP sites by using BLAT**^[10]^**. If the length of aligned sequence was shorter than 101 bp, this SNP was considered as incredible and it was removed; if the extracted sequence was aligned with the assembly results several times, this SNP was considered locate in repeat region and it was removed. Blast, TRF and Repeatmask software was used to predict SNP in repeat regions. The credible SNP was obtained through filtering SNP located in repeat regions.

#### (2) InDel detection

With LASTZ**^[11][12]^** software, the reference and query sequence CAED were aligned to get the alignment result. Through a series of treatment with axt-correction,axt-Sort, axt-Best, the best alignment results were chosen and the InDel result was preliminarily obtained. One hundred fifty base pairs in the upstream and downstream of InDel site in the reference sequence were extracted and then aligned with the query reads. The alignment result was verified with BWA**^[13]^** and samtools.

### 2.5 Transcriptome sequencing and comparison

#### (1) Sequencing and filtering

Total RNAs of SWS and SWG were extracted by TIANGEN RNAprep pure Kit (Beijing, China). Then we used mRNA enrichment or rRNA dedivision method to dispose Total RNAs. The first step involved purifying the poly-A containing mRNA molecules using poly-T oligo-attached magnetic beads, and then mRNA was fragmented into small pieces using divalent cations under elevated temperature. The cleaved RNA fragments were copied into first strand cDNA using reverse transcriptase and random primers. This was followed by second strand cDNA synthesis using DNA Polymerase I and RNase H. These cDNA fragments then had the addition of a single ‘A’ base and subsequent ligation of the adapter. The products were purified and enriched with PCR amplification. We quantified the PCR yield by Qubit and pooled samples together to make a single strand DNA circle (ssDNA circle), which resulted in the final library. We used internal software SOAPnuke to filter reads, followed as: 1) remove reads with adaptors; 2) remove reads in which unknown bases (N) are more than 10%; 3) remove low quality reads (we defined the low quality read as the percentage of base which quality was lesser than 15 was greater than 50% in a read).

#### (2) Gene expression value statistics

We mapped clean reads to the reference strain using HISAT(Version 2.0.4) and Bowtie2 (Version 2.2.5)**^[14]^**, and then calculated gene expression level with RSEM**^[15]^**. Cluster analysis of gene expression was performed using the software Cluster**^[16,17]^** (Version 3.0) and the results of cluster analysis were displayed with javaTreeview**^[18]^**.

#### (3) Differential gene expression analysis

Differential gene expression was analyzed using the DESeq method which based on the Poisson distribution and performed as described by Wang**^[19]^**. *P*-values were corrected to *Q*-values as the strategy previously described**^[20]^** to improve the accuracy of differentially expressed genes (DEGs). Genes with a Fold Change ≥ 2 and *Q*-value ≤ 0.001 in two different samples were defined as DEGs, and then the identified DEGs were enriched and clustered according to GO and KEGG database.

### 2.6 Proteomic analysis

#### (1) Polypeptide disposition

To explore the quantitative proteomics and examine the difference of protein profiles of these samples, we used high-throughput technology based on iTRAQ combined with two-dimensional liquid chromatography-tandem mass spectrometry (2D-LC-MS/MS) performed as previously described**^[21]^**. After extracted, quantified, SDS-polyacrylamide gel electrophoresis and trypsin digestion, the protein samples were processed into polypeptide and labeled with iTRAQ reagents. The polypeptide was separated by liquid phase separation and freeze-dried by liquid phase system (SHIMADZU LC-20AB) and then separated by nanoliter liquid chromatography (SHIMADZU LC-20AD). After liquid-phase separation, the peptides were ionized by nanoESI and entered into the series mass spectrometer Q-Exactive (Thermo Fisher Scientific, San Jose, CA) for DDA (data-dependent acquisition) mode detection and obtained raw data finally.

#### (2) iTRAQ quantification

The raw data were compared with the corresponding database by protein identification software Mascot to search and identify. To determine whether the data was qualified, the credible protein identification results were obtained. We used IQuant**^[22]^** software to quantify iTRAQ data, which integrated the Mascot Percolator**^[23]^** algorithm, so as to improve the identification rate of the result. Firstly, 1% FDR was filtered at the spectrum/peptide segment level (psm-level FDR ≤ 0.01) to get the spectrum chart of significance identification and the list of peptide segments. Then, based on the parsimony principle, the peptide fragments were used for protein assembly and produced a series of protein groups. In order to control the false positive rate of the Protein, the process was filtered again at the protein level by 1% FDR (protein-level FDR ≤ 0.01). The strategy used for this process was picked protein FDR**^[24]^**. In conclusion, we used iTRAQ quantitative analysis to screen out the significantly different proteins that we cared more about from the results. Finally, GO, Pathway functional annotations and GO, Pathway enrichment analysis of differentially expressed proteins (DEPs) were carried out.

## 3. Result

### 3.1 Phenotypic characteristics

#### (1) Morphology

The three strains showed no significant difference under the optical microscope, but when SEM was used to monitor single cell morphology, it was found there were obvious cracks on the cell surface of SWO and SWG, but no cracks appeared in SWS (Fig. 1).

**Figure 1.**
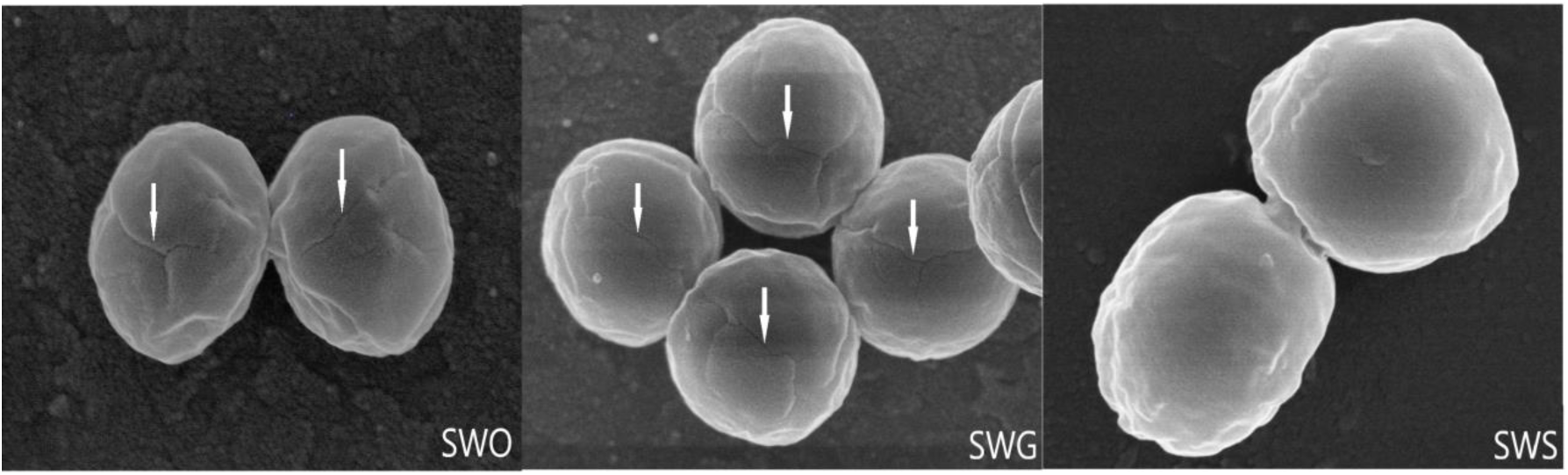
SEM morphology of the strains. Obvious cracks (white arrows) were found on the surface of SWO and SWG, while no cracks were found on the surface of SWS.

#### (2) Antibiotic susceptibility test

The results of the disk diffusion assays revealed that the three strains were sensitive to all tested antibiotic disks and the result showed no obvious difference among them (Table.1).

**Table 1.**
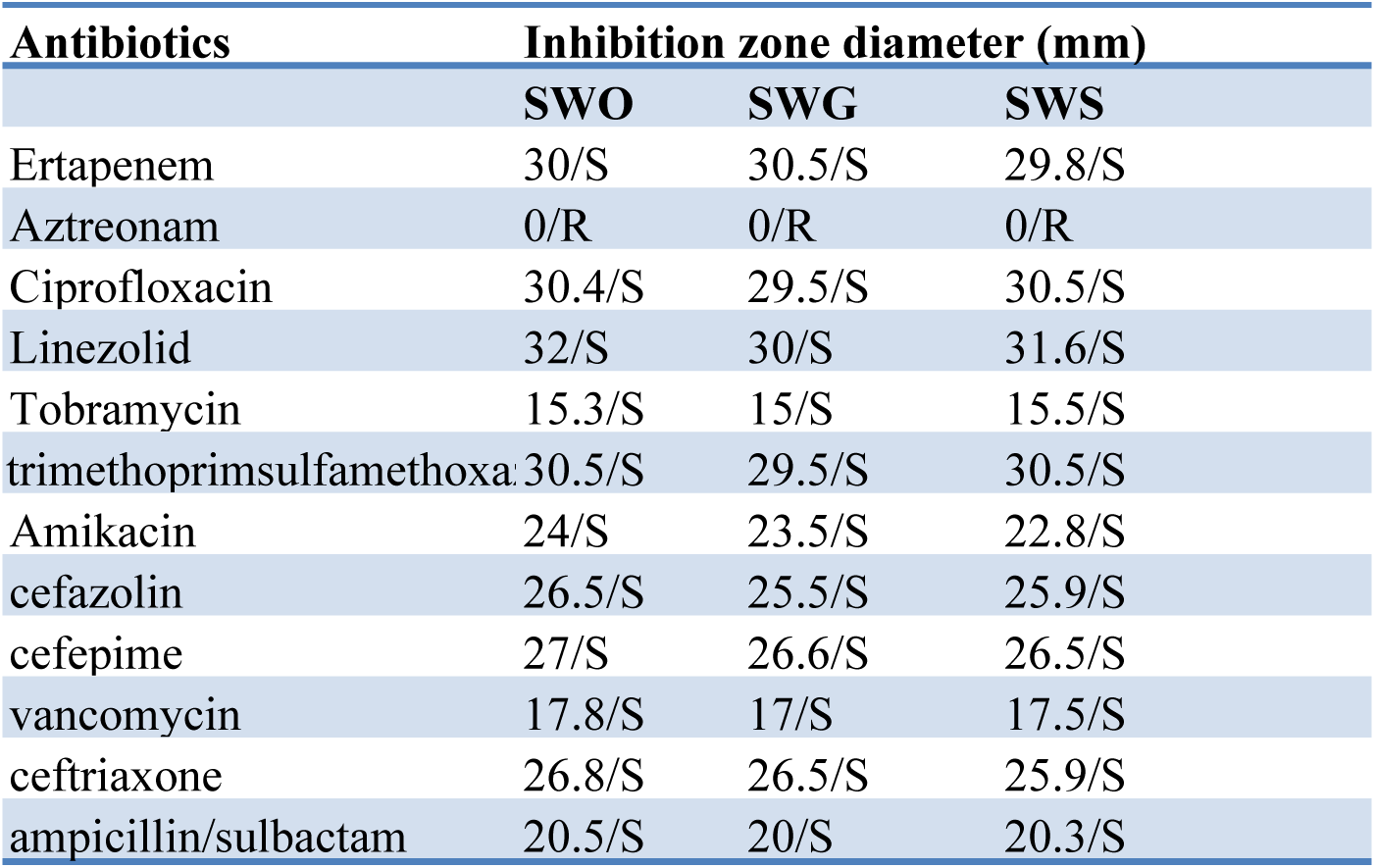
Antibiotic susceptibility analysis of the three strains

#### (3) Chemical sensitivity and carbon source utilization assay

Chemical sensitivity assay found that SWS displayed increased tolerance to potassium tellunite (PT), niaproof4 (N4), tetrazolium violet (TV) and sodium butyrate (SB), compared to SWG and SWO (*P* < 0.05) (Fig. 2). However, there were no significant difference between SWO and SWG. Carbon source utilization assays showed there was no significant difference among the three strains.

**Figure 2.**
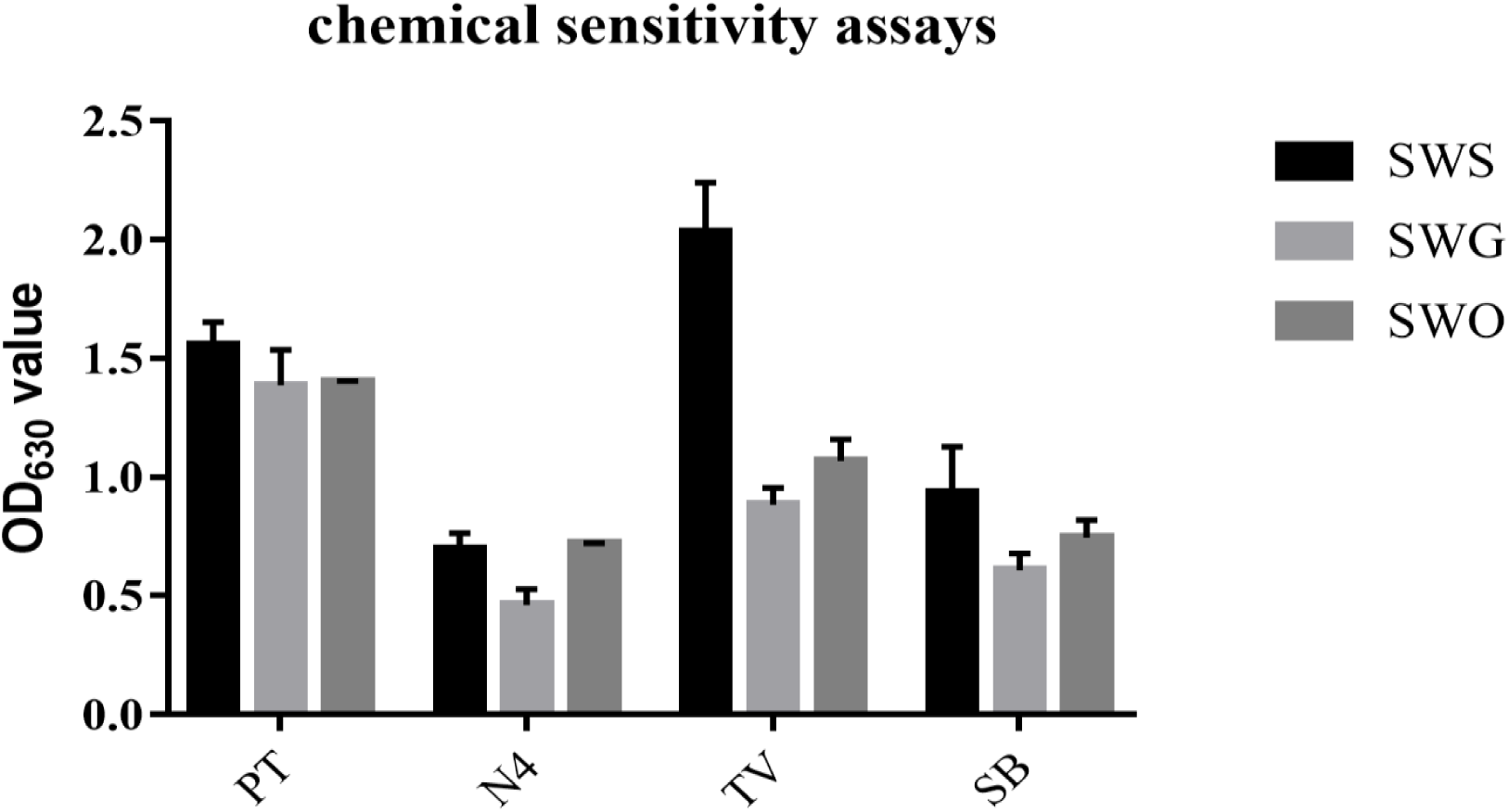
Chemical sensitivity assays of the three strains. SWO, SWG and SWS strain were incubated in Biolog GENIII MicroPlate at 37°C for 24 hr with LB medium and then tested carbon source utilization and chemical sensitivity ability of three strains. X-axis: potassium tellunite (PT), niaproof4 (N4), tetrazolium violet (TV) and sodium butyrate (SB)

#### (4) Growth rate assay

Some small difference was observed in the three strains, such as the growth rate and maximum OD_630_ values, but the difference was not significant (Fig. 3).

**Figure 3.**
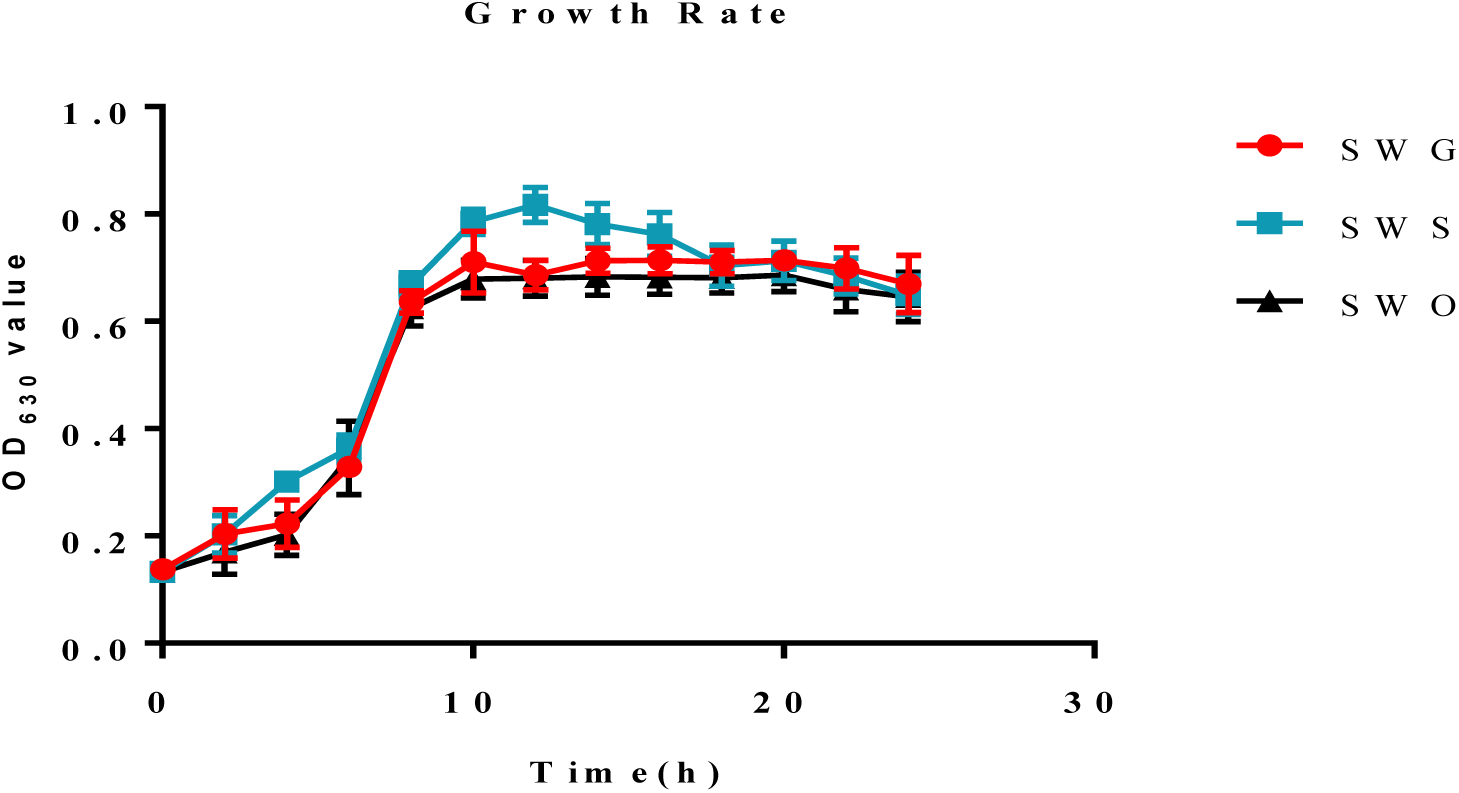
Growth rates of the three strains in LB medium. SWO,SWG and SWS strain were cultured in 20 ml LB medium at 37°C for 24 hr with agitation. OD_630_ was measured every 2 for 24 hr.

#### (5) Biofilm assay

The biofilm formation ability of bacteria is considered to be an significant phenotype because it is related to host colonization, antibiotic resistance, and environmental persistence, etc**^[25-26]^**. Therefore we performed biofilm test to estimate the biofilm formation ability of the three *S. warneri* strains. The result found that SWS showed the strongest biofilm formation ability and SWG and SWO hardly had this ability (*P* < 0.05) (Fig. 4).

**Figure 4.**
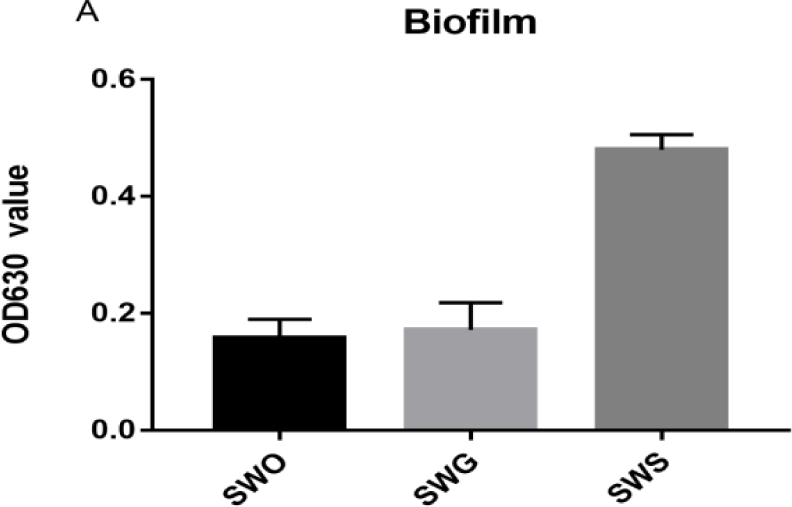
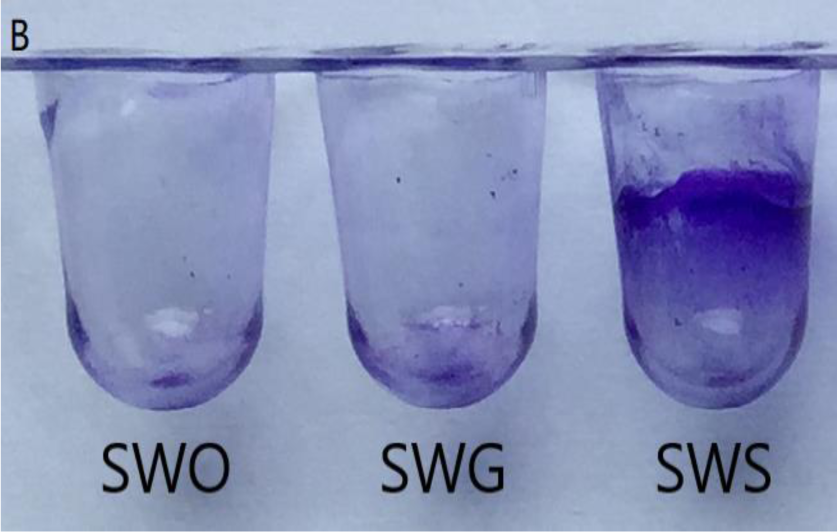
Analysis of biofilm formation ability via crystal violet staining. The three strains were cultured in 96-well polystyrene microtiter plates at 37°C for 24 hr. Biofilm formation ability was measured by determining OD_560_ of crystal violet.

### 3.2 Genomic sequencing, assembly and annotation

We made a complete genetic prediction and identification for SWO. After SWO complete genome were assembled, the results were used as the reference for comparative genomic analysis of SWS and SWG. Using Glimmer software**^[27]^** we identified 2,486 genes with a total length of 2,183,598 bp, which consisted of 84.88% of the genome and accession number of SWO is CP033098-CP033101. In addition, 5,745 bp of minisatellite DNA and 213 bp of mcrosatellite DNA sequences were identified, which composed of 22.33% and 0.83% of the genome, respectively.

All genes were annotated against the popular functional databases, including 65.32% of the genes into the GO database**^[28]^**, 75.98% of the genes into COG database**^[29]^**, 61.7% of the genes into KEGG**^[30]^**, 15.08% genes into NOG database and 99.07% of the genes into the NR database. Moreover, 16 genes were identified in the CAZY (Carbohydrate-Active enzymes) database**^[31]^**, 188 genes in the PHI-base(Pathogen Host Interaction) database**^[32]^**, 15 genes in the ARDB (Antibiotic Resistance Genes Database) database and 122 genes in VFDB (Virulence Factors Database) **^[33]^**. The genome map of SWO was shown in Fig. 5.

**Figure 5.**
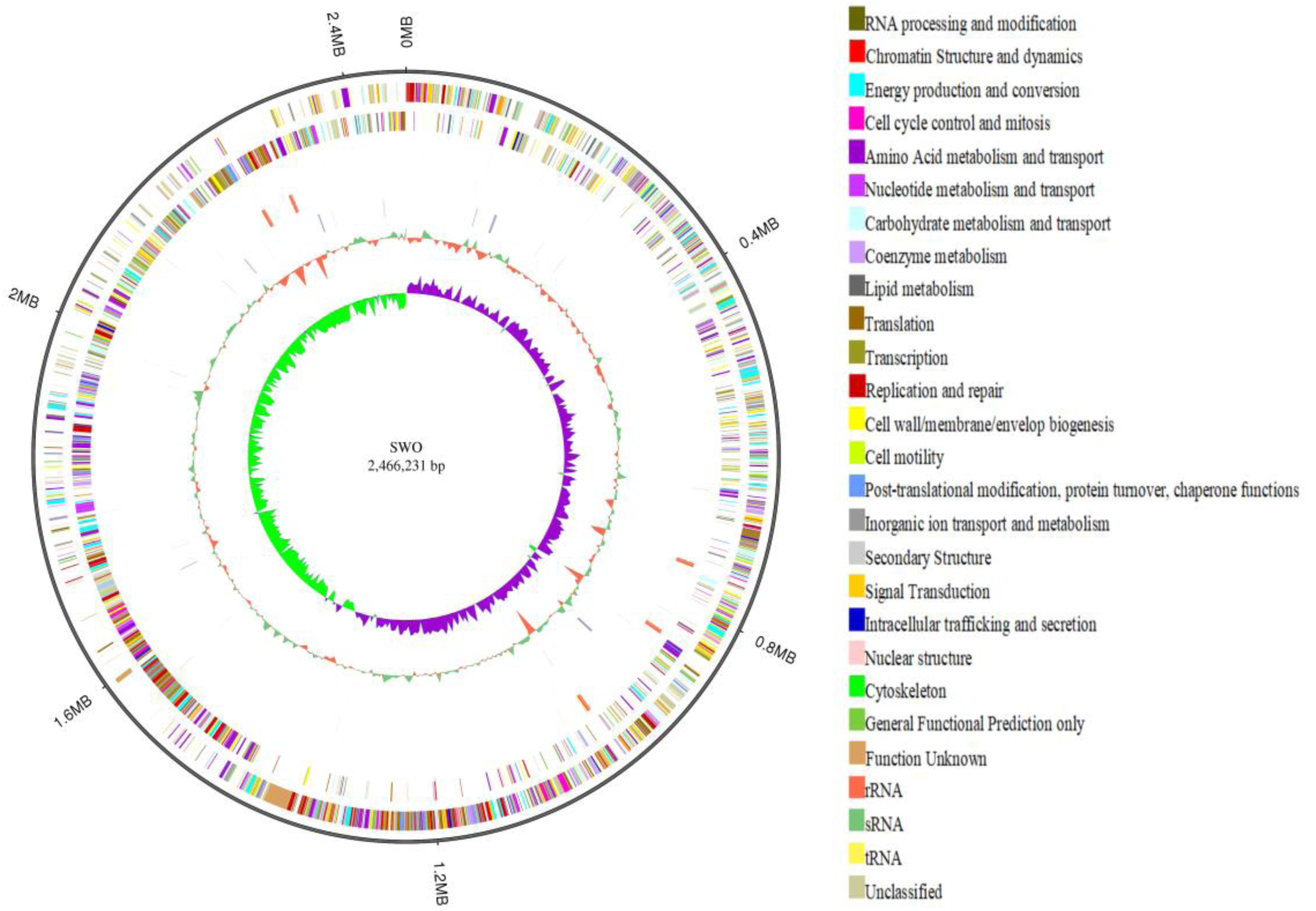
Genome map of SWO (ncRNA, COG annotation, GC content and GC skew). From outer to inner, the 1^st^ circle shows the genome size; the 2^nd^ circle shows the COG function of forward strand gene and each color represents a function classification; the 3^rd^ circle shows the function of reverse Strand Gene; the 4^th^ circle shows the ncRNA results of forward strand containing tRNA, rRNA and sRNA; the 5^th^ circle shows the ncRNA results of reverse strand; the 6^th^ circle shows repeat sequences; the 7^th^ circle shows the GC content and the 8th circle shows the GC skew ((G-C)/(G + C), green > 0, purple < 0).

### 3.3 Comparative genomic analysis

SWO was used as the reference strain to detected variations (SNPs and InDels) of SWS and SWG by using comparative genomics sequencing (Table. 2). The accession number for re-sequencing data of the SWS is SAMN10255186; SWG is SAMN10250545. Subsequently we conducted structure variations analysis of SWS and SWG compared to SWO.

**Table 2.** Statistics of Comparative genome sequencing.

| Sample | SWS | SWG |
| --- | --- | --- |
| **Raw read statistics** |  |  |
| Total reads | 2685888 | 2776954 |
| Sequencing depth (X) | 132 | 133 |
| Genome coverage (%) | 100% | 100% |
| **Assembly statistics** |  |  |
| Chromosome size | 2524548 | 2518925 |
| No. of scaffolds | 22 | 17 |
| Largest scaffold length | 687388 | 1265212 |
| N50 scaffold length | 318178 | 1265212 |
| G+C content (%) | 32.50% | 32.50% |

The raw variation sites were identified and then filtered with strict standards to detect potential InDels. After a series of filtering conditions, we found 15 InDels between SWS and SWG, including 9 InDels in intergenic regions and 6 in a coding region. Four were unique to SWS, 5 were unique to SWG, and all 9 were located in the intergenic regions. The CDS InDel was identified in SWOGL000595, SWOGL002370 and SWOGL002369, which is annotated as domain-containing protein.

### 3.4 Transcriptomic analysis

The accession numbers for transcriptomic data of SWS and SWG were SAMN10255187 and SAMN10255193, respectively. The average output of SWS and SWG was respectively 21.87 and 21.86 M. A total of 2271 and 2274 genes were expressed in SWS and SWG, including 2270 genes identified in both of the two strains. Analysis found both up-regulated and down-regulated genes were identified. Approximately 7 genes were up-regulated and 16 genes were down-regulated in SWS (Fig. 6). Simultaneously we found that the down-regulated genes significantly outnumbered up-regulated genes, suggesting that gene expression and metabolism were inhibited in SWS.

**Figure 6.**
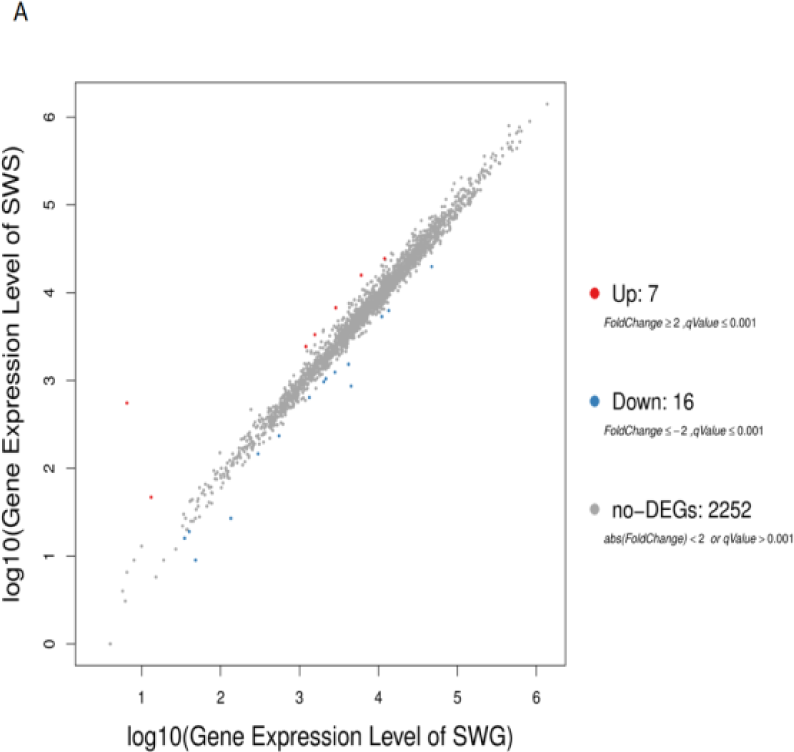
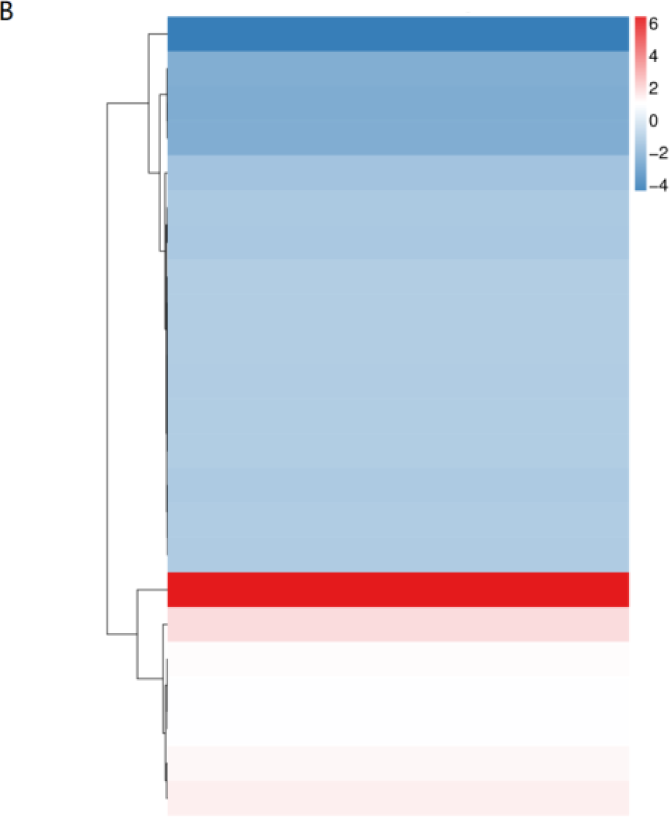
Differential transcriptomic expression **A.** These diagrams represent the number of genes that are specifically expressed in SWS and SWG. **B.** Hierarchical Clustering of DEGs. The X-axis represents the difference comparison for cluster analysis, and the Y-axis represents the difference gene. Color represents the difference multiple converted by log2.

We detected that there were not many gene changes in the two strains, suggesting that *S. warneri* was relatively stable under the stimulation of the external environment. In other words, it had a relatively strong resistance to the external environment.

DEGs were enriched and clustered according to GO and KEGG analysis. For GO analysis, the transcriptome of SWS was characterized by regulation of 4 DEGs involved in transporter activity, sub-function of molecular function, which was statistically significant. The up-regulated and down-regulated genes were summed and were compared with unchanged genes. We found that this function was significantly down-regulated by 3 genes and up-regulated by 1 genes (*P* < 0.05) (Table. 3).

**Table 3.**
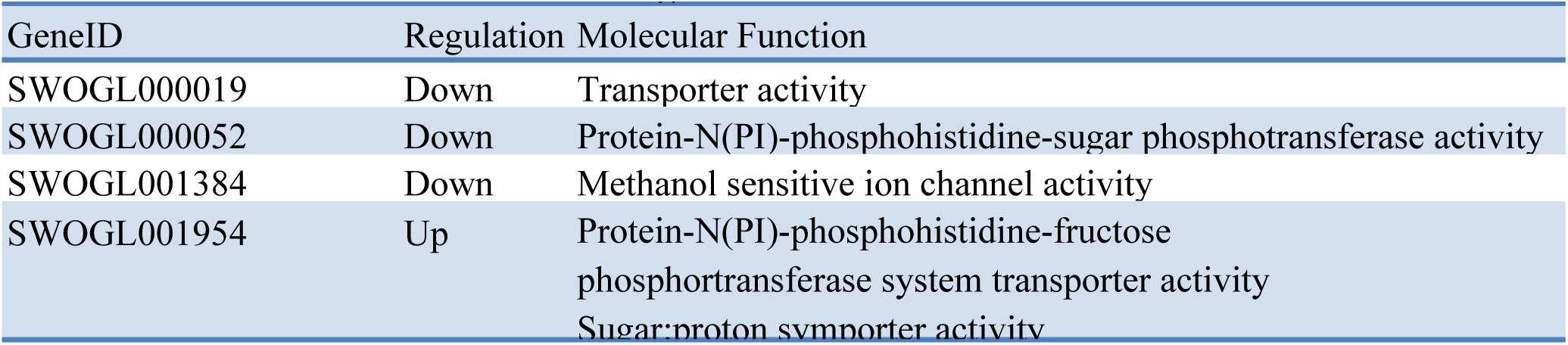
GO Functional annotation and regulation of 4 DEGs

Using the KEGG orthology based annotation system to identify metabolic pathways, we identified 2 DEGs (SWOGL001955, SWOGL001954) in fructose and mannose metabolism was statistically significant (*P* < 0.05) (Table. 4). The gene SWOGL001954 was also enriched to other 3 categories of KEGG. Different from the GO, the results of KEGG were all up-regulated.

**Table 4.**
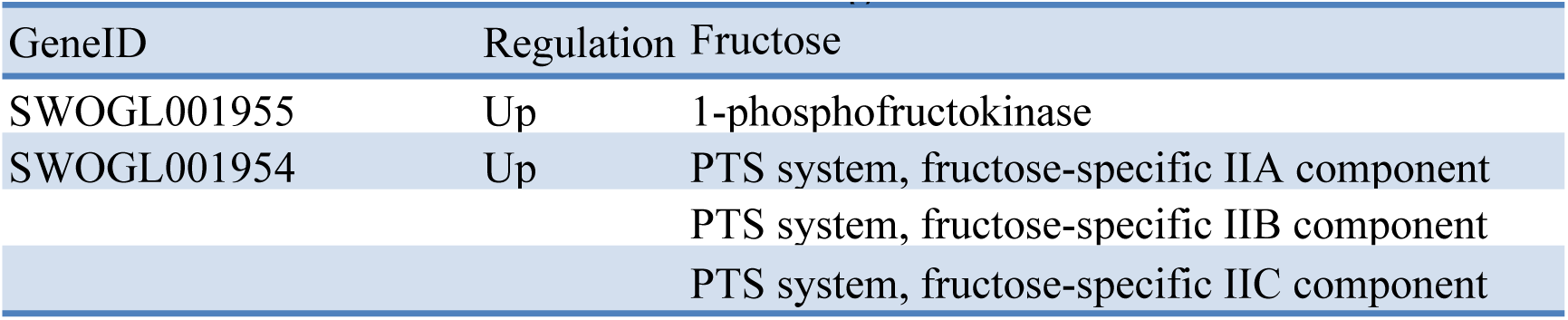
KEGG Functional annotation and regulation of 4 DEGs

### 3.5 Comparative proteomic analysis

Relatively quantitative analysis shows that 120 DEPs were identified, including 72 down-regulated and 48 up-regulated proteins (Fig.7).

**Figure 7.**
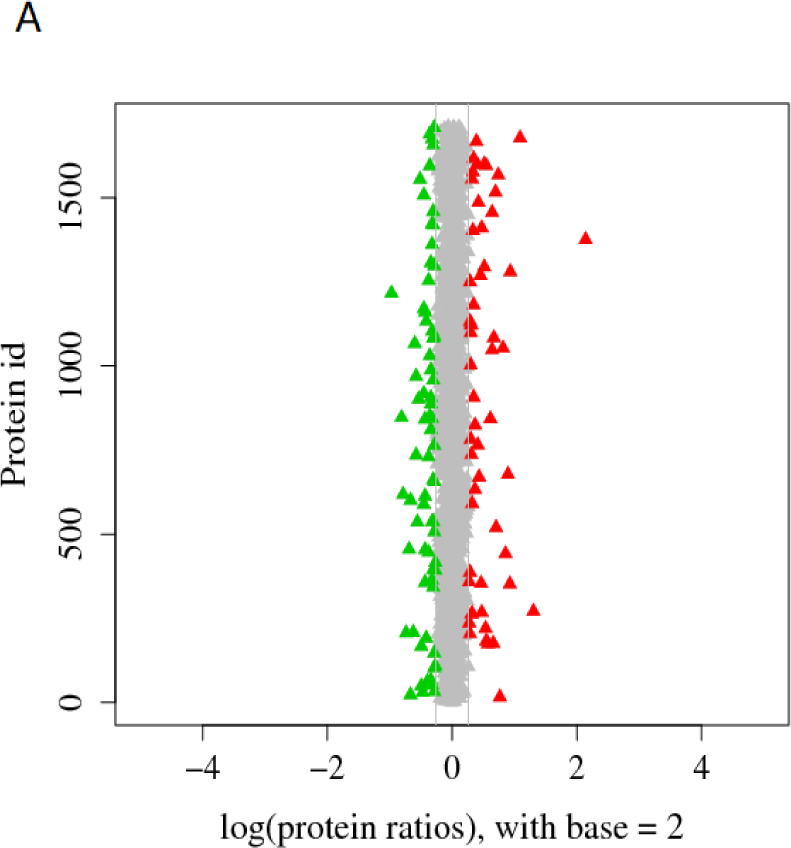
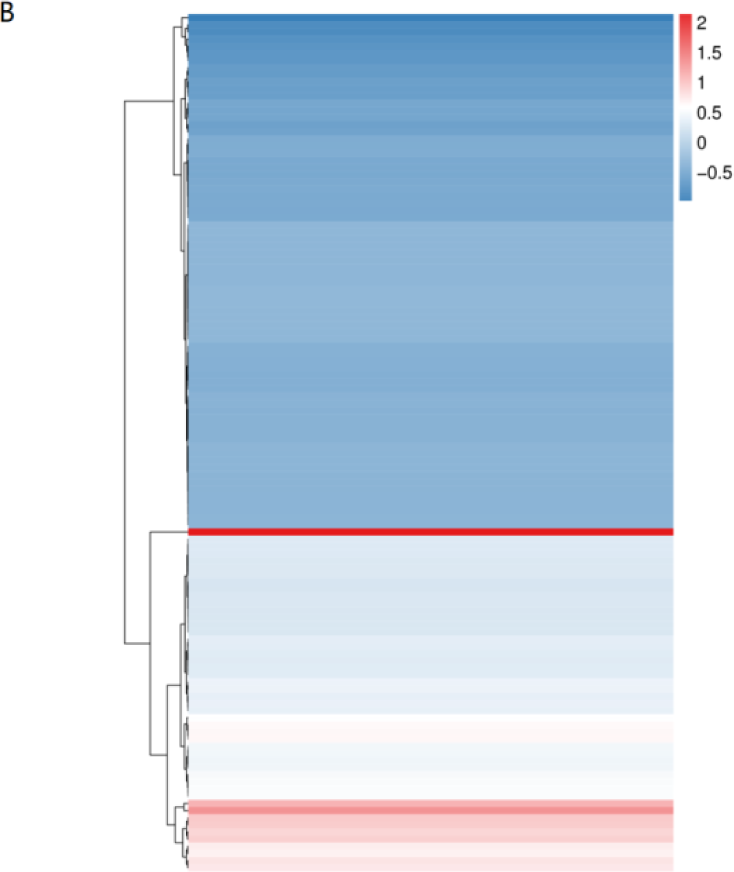
Differential proteomic expression. **A.** Protein ratio distribution of SWS and SWG. **B.** Hierarchical Clustering of DEPs.

Subsequently, DEPs were enriched and clustered according to GO and KEGG analysis. As to GO function category, it was clear that the expression of proteins involved in functions such as membrane, integral component of membrane, intrinsic component of membrane, cell periphery, membrane was statistically significant (*P* < 0.05). The up-regulated and down-regulated proteins were summed and were compared with unchanged proteins. The down-regulation of DEPs in GO were significant, and the proteins enriched to biological process and molecular function were basically down-regulated. With respect to KEGG functions, significant difference was found in the following pathways: cellular community-prokaryotes, carbohydrate metabolism and energy metabolism (*P* < 0.05) (Fig. 8A). Different from the transcriptomes, the KEGG of DEPs showed down-regulated proteins (Fig.8 B), including carbohydrate metabolism, and energy metabolism.

**Figure 8.**
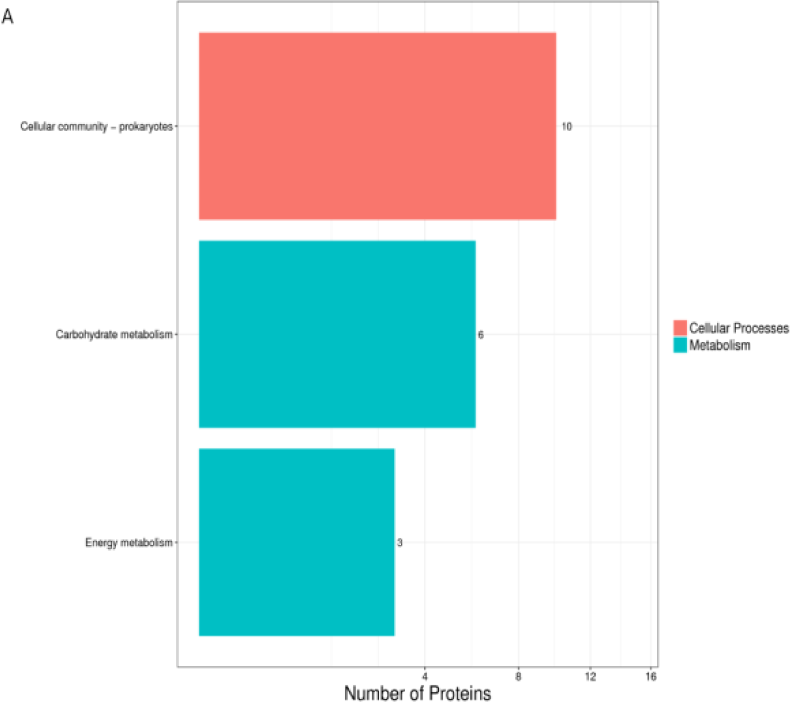
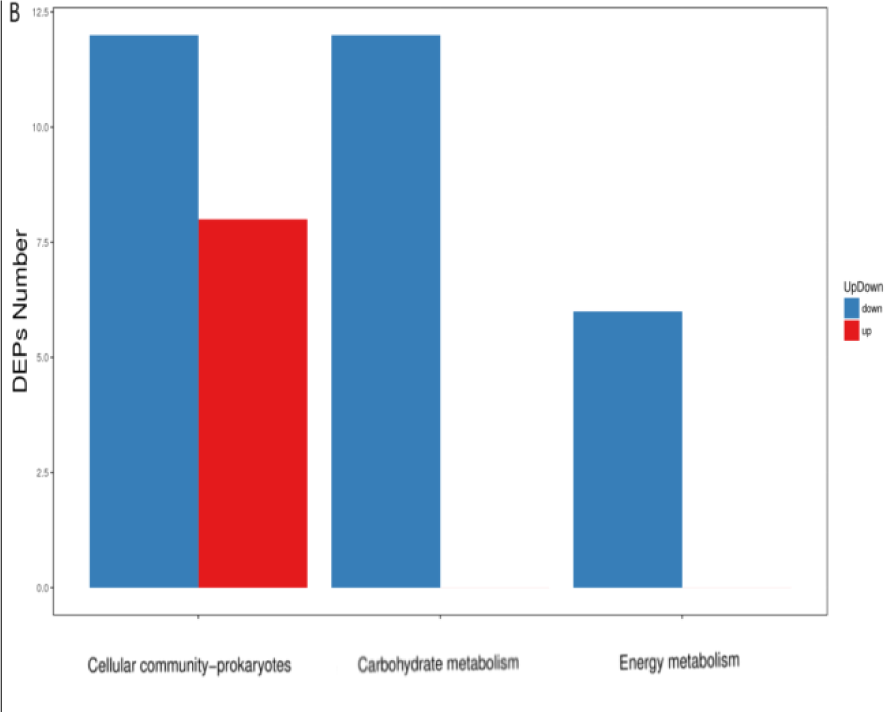
Differential proteomic analysis. **A:** Clustered DEPs in KEGG Pathway of SWS. The x-axis represents the number of the proteins corresponding to the GO functions. The y-axis represents KEGG Pathway functions. **B**: The regulation condition of DEPs enriched in KEGG Pathway of SWS. The x-axis represents the number of the proteins corresponding to the KEGG Pathway functions. The y-axis represents KEGG Pathway functions.

## 4. Discussion

The effect of space environment on microorganisms has been studied a lot, but the reports on *Staphylococcus* were less. In addition, the effect of long-term flight on bacteria has not been studied. A long-term space flight increases the risk of causing gene mutation of bacteria by the extended exposure to microgravity and strong high-energy radiation environment. In this study, we took advantage of this rare opportunity to research these two issues. *S. warneri* was carried as normal bacterial flora by the skin mucous membrane of the astronaut from Shenzhou-10. SWS had experienced of two flight processes, 15 and 64 days, respectively (see the description in the method for details). Some limited but significant changes between SWS/SWG/SWO were found at the genetic level. This gave us 2 points of suggestion.

1) Changes caused by space environment are mostly the genes related to resistance and adaptation to external pressure and stimulation. This can be demonstrated by the changes in the phenotype of SWS, including cell walls, biofilms and chemical sensitivity. The main component of *Staphylococcus* cell wall is peptidoglycan, and the mechanical strength of the cell wall of *S. warneri* basically depends on the existence of peptidoglycan. Thus the cell wall of *S.warneri* is closely related to the metabolic pathway of sugars. As an important feature of *Staphylococcus*, biofilm is defined as a sessile microbial community in which cells are attached to a surface or to other cells and embedded in a protective extracellular polymeric matrix. This mode of growth exhibits altered physiologies with respect to gene expression and protein production**^[35-37]^**. A major constituent of the biofilm matrix is polysaccharide intercellular adhesin (PIA) and the formation depends on phosphotransferase (PTS) system, a distinct method used by bacteria for sugar uptake. In the system the source of energy is from phosphoenolpyruvate (PEP), which involved in transporting many sugars into bacteria, including glucose, mannose, fructose and cellobiose. Although there is a general down-regulated trend in gene expression and protein products in SWS, there is still a significant up-regulation of PTS system in glucose metabolism. There are 4 genes involved in glucose metabolism, among which 3 were up-regulated and 1 was down-regulated. The up-regulation of gene SWOGL001954 plays an important role in the expression of PTS system. According to GO and KEGG database annotation, SWOGL001954 not only participates in the composition of the fructose-specific IIA/IIB/IIC components, but also takes part in the transport process of these complexes in PTS system. This indicates that the functions of SWS in manufacturing, utilization and transport of glucose metabolism associated with PTS system have been significantly enhanced. Therefore, it is believe that changes in biofilm and cell wall are directly related to PTS system enhancement of SWS. Peng**^[38]^** found PTS system in *Enterococcus fecal* helped to increase the bacteria’s resistance to stress in the environment. Therefore, it is reasonable to think that the decreased sensitivity of SWS to chemical stimulation is also related to the enhancement of PTS expression. Its chemical resistance might be achieved through a series of changes in biofilms and cell walls. DEPs enriched in KEGG showed a decrease in glucose metabolism, and we found only a correlation with PTS is lactose/cellobiose transporter subunit IIA (protein ID is WP_002450492.1). This protein is an enzyme with regulation function. For example, at low level of glucose concentrations the enzyme accumulates this activates membrane-bound adenylate cyclase. When the level of glucose concentration is high, the enzyme is mostly dephosphorylated and inhibits adenylate cyclase, glycerol kinase, lactose permease, and maltose permease. Therefore this protein is a bidirectional regulator in PTS. All the other DEPs down-regulated in glucose metabolism of KEGG function were related to galactose metabolism, which were independent of PTS. Galactose can be used for prevention of the biofilm formation by targeting activity of autoinducer 2 (AI-2), a quorum sensing molecule involving in virulence and biofilm formation^[39]^. This change confirms that the thickening of biofilm may related to the down-regulation of galactose metabolism.

2) The results of transcriptome and proteomics indicate that SWS is in a relatively low metabolic state. Therefore, it is reasonable to believe that SWS can increase its resistance to external environmental stimuli by lowering its own metabolism, and this change has indirectly led to even after a long space flight, the influence of *Staphylococci* on space environment changes is still relatively small, consistent with the results of Guo**[34]**. In addition, some results of other studies on the resistance of *Staphylococci* in the space environment were inconsistent. Tixador**^[40]^** found that the lowest inhibitory concentrations of benzocillin, erythromycin and chloramphenicol on *S. aureus* were increased. Similar changes were observed in *S. epidermidis*, the most abundant *Staphylococcus* in the space environment**^[41]^..**The study of fajardo-cavazos P**^[42]^** found that the mutation frequency of *rpoB* gene encoding rifampicin-resistant protein in *S. epidermidis* in space flight group was much higher than that of the ground group and the simulated gravity group. However, Rosado**^[43]^** found that the antibiotic sensitivity of *S. aureus* growing under simulated microgravity conditions was not significantly different from that of the control group. From the above, we speculate that different environments lead to different directional mutations in bacteria. The pressure of microgravity, radiation and other factors in the space environment to the bacteria often exists on the level of physical structure change and chemical damage. Therefore, if the bacteria itself are not pathogenic or opportunistic pathogens like *S. warneri*, changes in environmental adaptation, physical resistance and cell repair should be more obvious or even become stable mutations, while changes in such aspects as drug resistance and virulence should not be significant because there is no threat of antibiotics, and our results confirm our suspicions.

At the same time, we also noticed that the control group SWG also had limited changes in both phenotype and genetic level compared to SWO after dozens of days cultivation. Therefore, it is reasonable to assume that the mutation in the space environment of *S. warneri* can be inherited stably after brought back to the ground. If SWO isolated directly from space can be compared with the primitive *S. warneri* on the ground to prove that the mutation is indeed caused by the influence of space environment, our assumption should be more convincing. This is also worth further study.

## 5. Conclusion

In this study, the effects of long-term flight on bacterial mutation were studied using *S. warneri*. The results showed that PTS of *S.warneri* was up-regulated by long-term space stimulation, which resulted in a series of changes in cell wall, biofilm and chemical sensitivity, thus enhancing the resistance and adaptability of bacteria to the external environment. No other significant mutations were found in drug resistance. However, as an opportunistic pathogenic bacterium, the infection and sepsis caused by *S. warneri* are diseases that cannot be easily cured even in the ground environment with complete medical conditions. Once similar cases are found in the space capsule environment, it will be a disaster. Therefore, in the space capsule environment, we should not neglect any potentially pathogenic bacteria. How short-term human-bacterial symbiosis affects the human body and any changes in the biological characteristics of *Staphylococci* after long-term space flight are also worthy of exploring in the future.

